# Amplification-free, CRISPR-Cas9 Targeted Enrichment and SMRT Sequencing of Repeat-Expansion Disease Causative Genomic Regions

**DOI:** 10.1101/203919

**Authors:** Yu-Chih Tsai, David Greenberg, James Powell, Ida Höijer, Adam Ameur, Maya Strahl, Ethan Ellis, Inger Jonasson, Ricardo Mouro Pinto, Vanessa C. Wheeler, Melissa L. Smith, Ulf Gyllensten, Robert Sebra, Jonas Korlach, Tyson A. Clark

## Abstract

Targeted sequencing has proven to be an economical means of obtaining sequence information for one or more defined regions of a larger genome. However, most target enrichment methods require amplification. Some genomic regions, such as those with extreme GC content and repetitive sequences, are recalcitrant to faithful amplification. Yet, many human genetic disorders are caused by repeat expansions, including difficult to sequence tandem repeats.

We have developed a novel, amplification-free enrichment technique that employs the CRISPR-Cas9 system for specific targeting multiple genomic loci. This method, in conjunction with long reads generated through Single Molecule, Real-Time (SMRT) sequencing and unbiased coverage, enables enrichment and sequencing of complex genomic regions that cannot be investigated with other technologies. Using human genomic DNA samples, we demonstrate successful targeting of causative loci for Huntington’s disease (*HTT*; CAG repeat), Fragile X syndrome (*FMR1*; CGG repeat), amyotrophic lateral sclerosis (ALS) and frontotemporal dementia (*C9orf72*; GGGGCC repeat), and spinocerebellar ataxia type 10 (SCA10) (*ATXN10*; variable ATTCT repeat). The method, amenable to multiplexing across multiple genomic loci, uses an amplification-free approach that facilitates the isolation of hundreds of individual on-target molecules in a single SMRT Cell and accurate sequencing through long repeat stretches, regardless of extreme GC percent or sequence complexity content. Our novel targeted sequencing method opens new doors to genomic analyses independent of PCR amplification that will facilitate the study of repeat expansion disorders.

## Introduction

Eukaryotic genomes frequently contain regions that are not well addressed by commonly employed sequencing technologies (Ashley 2016). These include regions of extreme GC content, repetitive sequence, and sequence context that have reduced amplification efficiency. Most targeted sequencing approaches that typically rely on amplification cannot fully address these genomic regions (Mertes et al. 2011).

One such class of sequences are tandem repeat tracts that vary in length between different individuals, and that cause disease when expanded beyond certain thresholds. The repeats often become unstable as they get longer and can dramatically expand to hundreds or thousands of repeat units (Ashizawa et al. 1993; Veitch et al. 2007; Swami et al. 2009). There are more than 30 human disorders associated with repeat expansions including Fragile × syndrome, Huntington’s disease, myotonic dystrophy, amyotrophic lateral sclerosis/fronto-temporal dementia, and spinocerebellar ataxias (Orr and Zoghbi 2007; La Spada and Taylor 2010; Lopez Castel et al. 2010). Correct diagnosis and a more informed prognosis of these diseases relies upon accurate mutant repeat size estimates. In addition, interruptions of the repeat may modulate stability, disease heritability (Yrigollen et al. 2012) and disease phenotype (Matsuyama et al. 1999; Sakamoto et al. 2001; Charles et al. 2007; Braida et al. 2010; McFarland et al. 2014). Thus, it is important to obtain complete and accurate nucleotide-level resolution of these regions, in addition to repeat length.

Much work has been done to develop assays to measure repeat expansions (Warner et al. 1996; Adler et al. 2011; Bastepe and Xin 2015; Hayward et al. 2016). However, with respect to next generation sequencing (NGS) methods, these are some of the most challenging regions of the genome due to the potential for very long repeating sequence and, in some cases, very high GC content (Loomis et al. 2013; Ardui et al. 2017) which hinders amplification kinetics. These two factors also provide a significant challenge for PCR-based assays to faithfully reproduce the native genomic sequence during amplification. Furthermore, polymerase chain reaction (PCR) in patients heterozygous for a normal and large expansion allele can be problematic as only the normal allele may be amplified (Chakraborty et al. 2016) and polymorphisms surrounding the repeat region can lead to allele bias, dropout, or misinterpretation of results (Bastepe and Xin 2015). To address these limitations, we have developed a novel targeted sequencing approach that does not rely on amplification and employs Single Molecule, Real-Time (SMRT) sequencing that does not suffer from the read length and GC-bias shortcomings present in first and second-generation sequencing technologies (Loomis et al. 2013; Roberts et al. 2013; Ross et al. 2013; Shin et al. 2013; Chaisson et al. 2015). The method utilizes the site-specific endonuclease activity of the CRISPR-Cas9 system to target, enrich, and isolate specific DNA molecules. While Cas9 has become widely used for genome engineering (Cong et al. 2013; Mali et al. 2013), here we are simply using its properties as an RNA-guided DNA endonuclease (Jinek et al. 2012). CRISPR-Cas9 has been used previously for targeted sequencing (Shin et al. 2017), but not in conjunction with SMRT Sequencing which allows the sequencing of disease-associated repeats that are not covered using short-read sequencing technologies. We demonstrate the capabilities of the method by capturing multiple loci associated with repeat expansion disorders and sequencing of DNA samples with varying repeat lengths.

## Results

### Amplification-Free Enrichment Via Cas9 Targeting

An overview of the method is outlined in Figure 1. Briefly, SMRTbell libraries were generated from native genomic DNA digested with a combination of two restriction enzymes, EcoRI-HF and BamHI-HF (Travers et al. 2010). We then cut open the SMRTbell templates that contained our region of interest using Cas9 and a crRNA designed to be complementary to a sequence adjacent to the region of interest. A new hairpin adapter (capture adapter) was subsequently ligated to the digested templates and that adapter was used as a handle in MagBead capture to specifically enrich for the region of interest. Captured SMRTbell templates were sequenced using PacBio SMRT Sequencing (Eid et al. 2009). The unique properties of SMRT Sequencing allow for long reads which can span repeats many kilo-bases (kb) in length and provide unbiased coverage making it possible to sequence through the long stretches of high GC content present in some expanded repeats (Loomis et al. 2013; Roberts et al. 2013; Ross et al. 2013; Shin et al. 2013; Chaisson et al. 2015).

**Figure 1:**
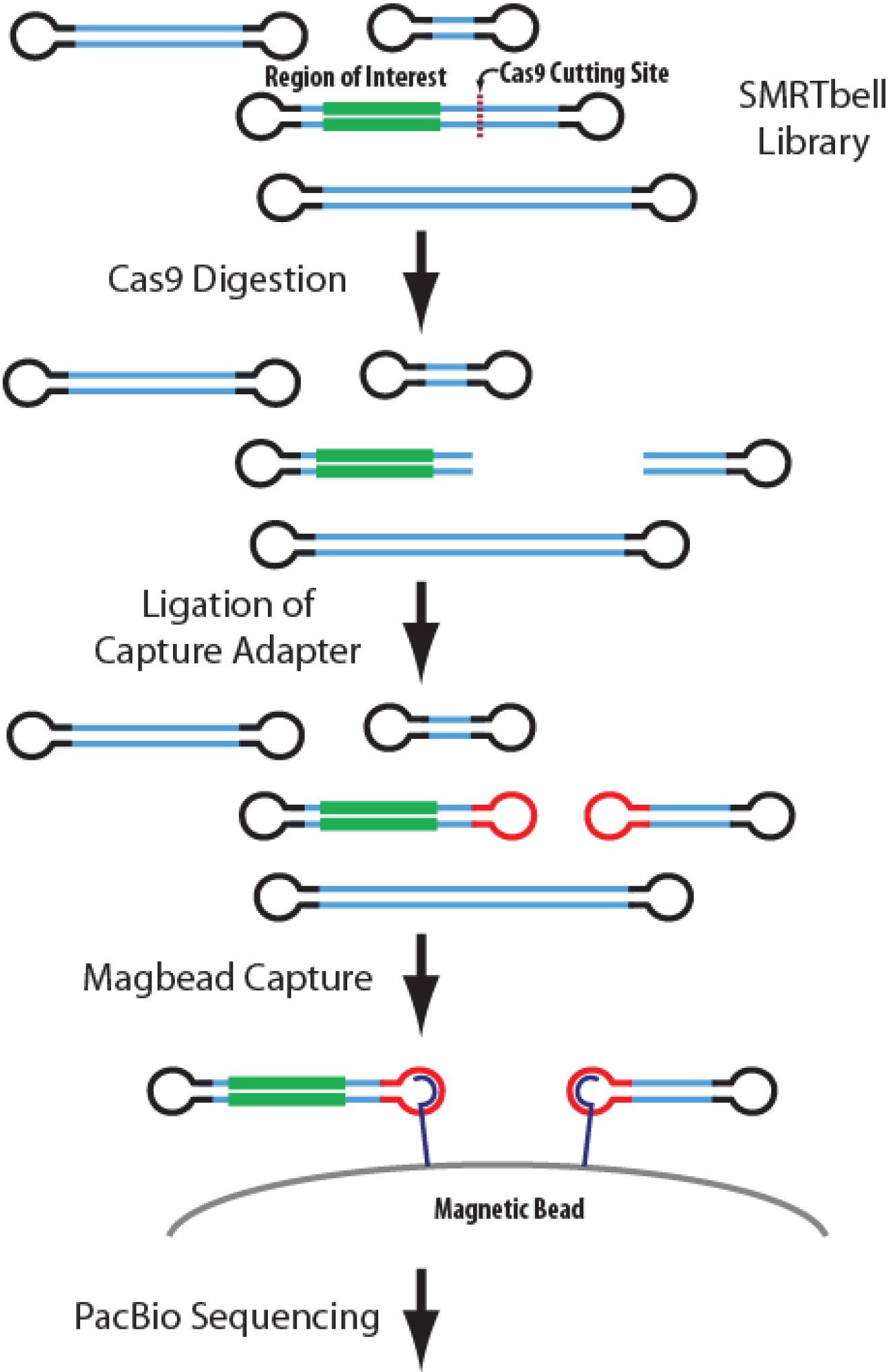
Overview of Method. An overview of the general concept of employing Cas9 for non-amplification based targeted enrichment. A standard SMRTbell library is created and a crRNA is designed adjacent to the region of interest (green). Digestion with Cas9 breaks open the SMRTbell molecules to enable ligation with a capture adapter. SMRTbell molecules that contain the capture adapter are enriched on magnetic beads and prepared for SMRT Sequencing on a PacBio RS II.

We designed Cas9 guide RNAs (crRNAs) for four targets with well-studied repeat expansions that are associated with human disease. These loci included *HTT*, *C9orf72*, *ATXN10*, and *FMR1*. Details of the repeat sequences and genomic locations are outlined in Table 1. The crRNAs can be combined during the enrichment and sequencing template preparation process, allowing for the simultaneous targeting of multiple loci from the same sample in a single experiment. The method was first tested on a control DNA sample extracted from HEK-293 cells. This sample was not expected to have expanded repeats in any of the four target loci. It should also be noted that this cell line has abnormal ploidy, with some regions of the genome having increased copy number. The four guide RNAs were designed to recognize a sequence adjacent to the repeat stretch of the four target loci (Supplemental Figure 1). After Cas9 digestion and ligation of the capture adapter, MagBead captured SMRTbell templates were subjected to SMRT Sequencing.

**Table 1:** Target Loci.

| Target | Associated Disease(s) | Chr | crRNA Coordinates (GRCh38) | Strand | Target Size | Repeat | Comments |
| --- | --- | --- | --- | --- | --- | --- | --- |
| *HTT* | Huntington's disease | Chr4 | 3075105-3075086 | - | 1125 bp | CAG | Common SNP (rs2857935) disrupts BamHI site, causing an upstream (1736 bp) BamHI site to be used. Coriell samples NA03620 and NA06903 have alleles with this SNP. |
| *C9orf72* | familial frontotemporal dementia (FTD) and amyotrophic lateral sclerosis (ALS) | Chr9 | 27573257-27573276 (a)<br>27572970-27572989 (b) | + | 1261 bp | CCCCGG | Genscript DNA sample has a SNP that adds a BamHI site decreasing the capture size to 1026 bp. |
| *ATXN10* | spinocerebellar ataxia Type 10 (SCA10) | Chr22 | 45794847-45794866 | + | 1019 bp | variable<br>ATTCT |  |
| *FMR1* | Fragile X and Fragile X-associated Tremor/Ataxia Syndrome (FXTAS) | ChrX | 147911587-147911606 | + | 1013 bp | CGG |  |

Supplemental Figure 2 shows the read length distribution for this sequencing run. The average polymerase read is greater than 16 kb in length. We designed the size of the captured regions to be approximately 1 kb according to the reference genome (hg19), however, the size of the captured molecules will increase for samples with expanded repeats. Due to the circular topology of the SMRTbell template, one sequencing read will consist of multiple passes over a single DNA molecule (Travers et al. 2010) and this allows for generation of an intramolecular CCS reads. CCS reads were generated using the PacBio SMRTPortal software and mapped to the human genome reference (hg19). On average, intra-molecular consensus sequences were generated from reads containing >14 passes and have predicted consensus accuracies of ^~^99% (Supplemental Figure 2).

### Complexity Reduction Improves On-Target Rate

To further improve the on-target fraction, we employed a strategy to enzymatically digest a large fraction of the genome prior to Cas9 targeting, thereby effectively reducing the size of the genome from which the desired DNA fragments are isolated. Restriction enzymes that are not predicted to cut inside the captured regions of interest were selected (KpnI-HF, MfeI-HF, SpeI-HF, EcoRV-FH), and prior to making EcoRI/BamHI genomic SMRTbell libraries, the gDNA was digested with either two or four of these complexity-reduction restriction enzymes, in the presence of phosphatase. As a result, the unwanted DNA fragments become incompatible with SMRTbell adapter ligation and are subsequently removed by treating the final SMRTbell library with exonuclease. Based on *in silico* digestion of the human genome, two complexity reduction restriction enzymes are predicted to remove ^~^62% of the human genome, while four enzymes can eliminate ^~^78% of the human genome. The amount of complexity reduction is also reflected in the yield of SMRTbell library (Supplemental Table 1).

We captured the four targets from the HEK-293 control sample that had been pre-treated using no, two, or four complexity-reduction restriction enzymes. A constant mass (approximately 1 μg) of SMRTbell library was taken into the Cas9 targeting step. While the number of total sequencing reads per sample remained roughly constant at ^~^50,000, inclusion of the complexity-reduction step significantly increased the total number of captured molecules for each of the four targets (Figure 2 and Supplemental Table 1), thereby increasing the on-target fraction from ^~^2% to more than 9%. On-target fraction is calculated by dividing the total number of molecules that map to any of the four target loci by the total number of sequencing reads that map to the human genome (hg19). For all four targets, this equates to an enrichment factor of more than 64,000.

**Figure 2:**
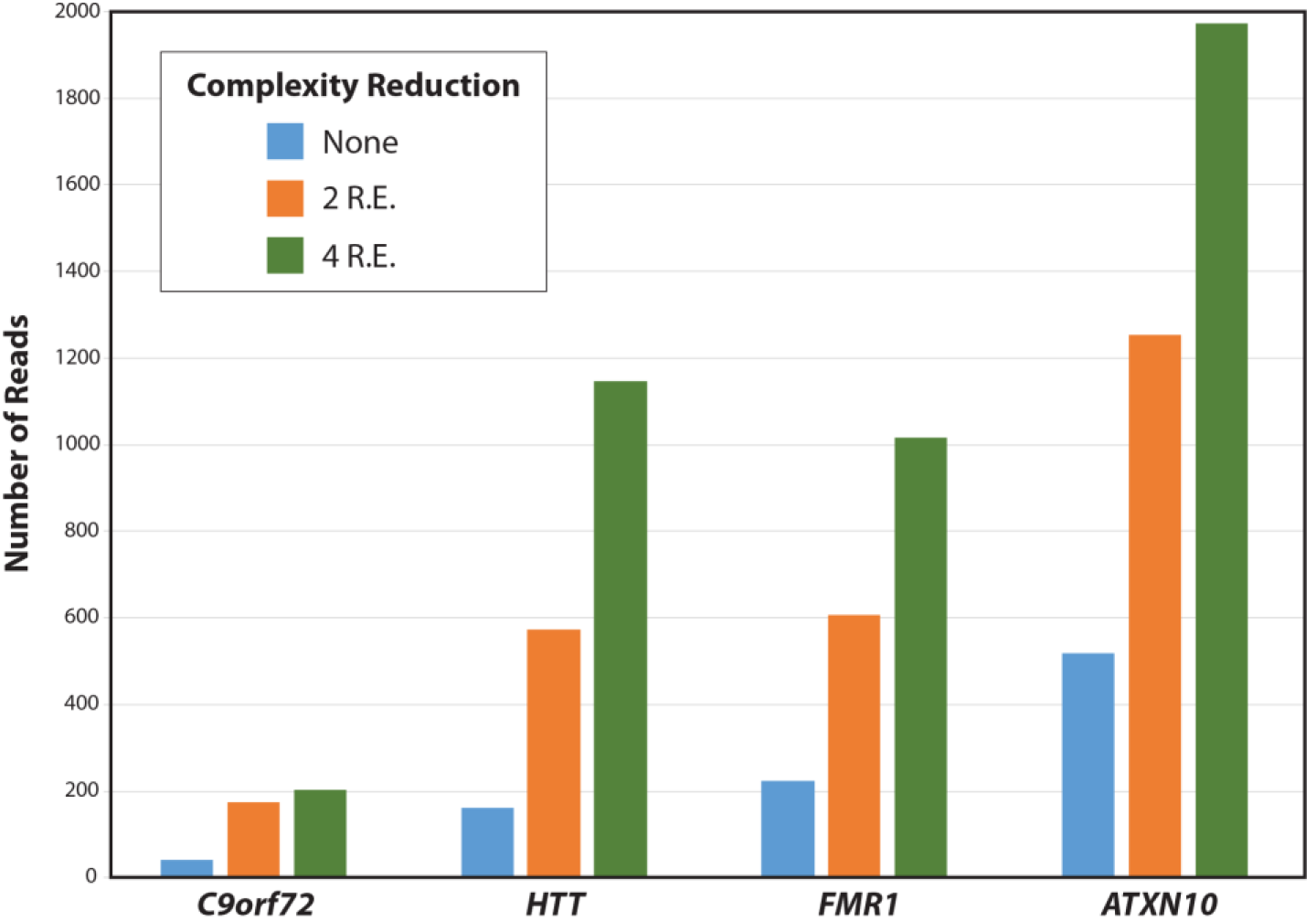
Complexity Reduction Via Restriction Enzyme Degradation Increases the Number of On-Target Reads. Number of CCS reads (individual molecules) is plotted for each of the four target regions (*C9orf72*, *HTT*, *FMR1*, and *ATXN10*). Colors represent the amount of complexity reduction employed (blue = none; orange = two restriction enzymes (KpnI-HF and MfeI-HF), green = four restriction enzymes (KpnI-HF, Mfe-HFI, SpeI-HF, EcoRV-HF)).

A coverage plot across the entire human genome organized by chromosome with the full set of four complexity-reduction enzymes is shown in Figure 3. Four of the distinct peaks are labeled and represent the four target loci. Other significant peaks (with more than 10-fold coverage) can be explained by off-target Cas9 digestion of instances in the human genome with sequence similarity to the designed guide RNAs, which is expected for 20-mer guide RNAs in a large genome (Supplemental Figure 3). The lower coverage of these off-targets may be explained by imperfect matches to guide RNAs. Interestingly, one off-target site on chromosome 9 is found only in the HEK-293 control DNA sample and not seen in other DNA samples used in this study. The HEK-293 DNA sample has a single nucleotide polymorphism (SNP) that increases the homology to the *ATXN10* crRNA (Supplemental Figure 3-C). Remaining off-target reads were randomly distributed across the genome and likely represent general background of the MagBead enrichment step.

**Figure 3:**
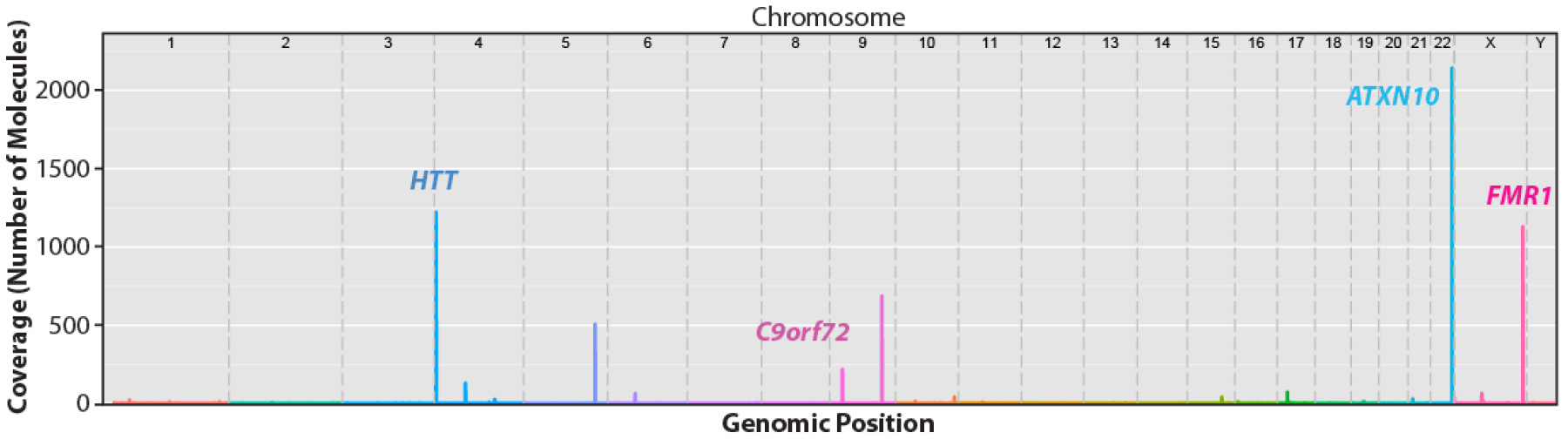
Coverage Across Genome. Maximum coverage within a window is plotted across the human genome reference (hg19) for a single capture experiment with four guide RNAs. Coverage is based upon the number of CCS reads which corresponds to the number of individual molecules captured and sequenced. The human genome reference is arranged by chromosome. Genomic positions of the four targets (*HTT*, *C9orf72*, *ATXN10*, and *FMR1*) are labeled.

The *C9orf72* locus consistently showed lower capture and sequencing efficiency as compared to the other three target loci. This is likely due to a SNP in the HEK-293 control DNA sample that introduces an internal BamHI site, decreasing the size of the captured molecule below the optimal range for efficient MagBead loading onto the SMRT Cell. The optimal fragment size is between 1-2 kb, though longer fragments up to approximately 10 kb will still load efficiently. For all subsequent experiments, we designed a new guide RNA (labeled “b” in Table 1) for the *C9orf72* locus that increases the size of the captured molecule from 742 bp to 1030 bp (Supplemental Figure 1). This new guide RNA moderately improved the number of on-target reads captured and sequenced for this locus. In addition, the efficiency of the Cas9 digestion may be low due to a suboptimal crRNA design. We do not believe the repeat content of the *C9orf72* locus is having significant impact because sequencing control samples from plasmids containing expanded GGGGCC hexanucleotide repeats did not show decreased read depth (data not shown).

### Capturing and Sequencing Expanded Repeats

To test the method on samples with expanded repeats associated with disease, we applied the method to genomic DNA samples from lymphoblastoid cells lines (LCL) generated from patients with repeat expansions in either the *HTT* (CAG repeat) or *FMR1* (CGG repeat) genes (see Materials and Methods for sample details). Results from two of these samples are shown in Figure 4. Sample NA03620 (Figure 4A) had previously been shown to contain an expansion in the *HTT* locus, with repeats of 18 and 60 (Maglione et al. 2010), with the longer allele being in the pathogenic range for Huntington’s disease. Using the same enrichment and sequencing protocol described above, we obtained CCS reads from more than 400 individual non-amplified molecules, with approximately equal sequencing coverage of each of the two alleles. Repeat lengths matched well with previous results, showing a bimodal distribution of repeat lengths with modes of 18 and 60 CAG repeats. The alleles can be segregated not only by CAG repeat length, but also by the length of a short stretch of CCG repeats just downstream of the CAG repeat (blue region in Figure 4A). The lengths of this polymorphic repeat show different distributions among normal and HD chromosomes, but do not contribute to Huntington’s disease onset age (Andrew et al. 1994). The distribution of CAG repeat lengths broadened for the expanded allele, reflecting mosaicism previously observed in HD patient tissues and cells (Kennedy et al. 2003; Veitch et al. 2007; Swami et al. 2009). The increased resolution of our method allows for the accurate characterization of the distribution of repeat sizes, which may further aid the study of somatic repeat instability (Swami et al. 2009; Lee et al. 2010). In sample NA03620, the normal allele with 18 CAG repeats has a fairly common single nucleotide polymorphism (SNP rs2857935) that disrupts the upstream BamHI site. The result is that an upstream BamHI site is used (Supplemental Figure 1) increasing the captured fragment size by 1736 bp.

**Figure 4:**
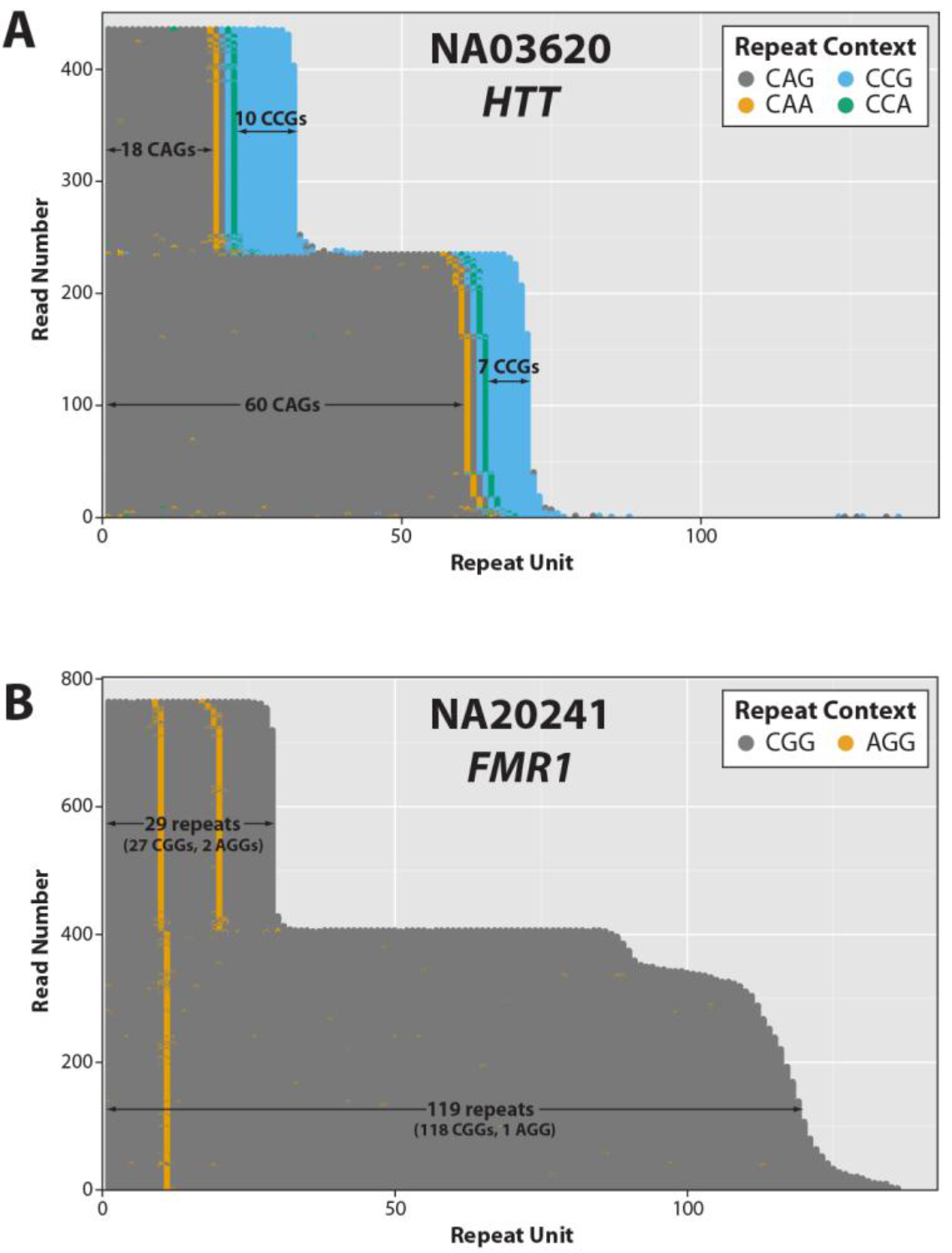
Visualization of Repeat Structures. Individual CCS reads are trimmed of flanking sequence to include only the relevant repeat region. Trimmed repeat sequences are sorted from shortest to longest and each individual molecule is represented by a series of colored dots on a horizontal line. Each dot represents a single repeat unit, color coded based on the repeat sequence. (A) *HTT* region in NA03620: the normal allele has a modal CAG repeat of 18, with 10 CCG, while the mutant allele has a modal CAG repeat of 60, with 7 CCGs. (B) *FMR1* region in NA20241: the normal allele has a modal total repeat count of 29, in which the CGG repeat is interrupted with 2 AGGs, while the mutant allele has a modal total repeat count of 119, in which the CGG repeat is interrupted with 1 AGG.

Sample NA20421 (Figure 4B) is derived from a patient with a pre-mutation length expanded CGG repeat in the *FMR1* gene. Repeat sizes were previously measured at 29 and 93-110, which was determined by testing in 9 independent laboratories using two methods each; however, neither of these methods provided a consensus repeat length on the expanded allele (Amos Wilson et al. 2008). We generated consensus sequences from nearly 800 individual molecules for the *FMR1* locus in this sample, relatively well distributed between the two alleles. The two alleles were segregated both by CGG repeat length and number, as well as the position of AGG interruptions (yellow stripe in Figure 4B). Interestingly, the expanded allele appears to have a bimodal distribution, with most molecules centered around 120 repeats, but a subset of molecules with closer to 90 repeat units. We obtained consensus sequences that fully span this range (Supplemental Figure 4). While mosaicism in expanded repeats is expected, this may explain the difficulty in accurately sizing the repeat lengths in previous studies.

The repeat length distributions of two more Coriell samples with *FMR1* expansions were determined (Figure 5). Coriell sample NA06903 is predicted to have CGG repeat lengths of 23 and 95. Our results were similar with modal repeat lengths of 24 and 93. Based on previous studies, the NA07537 sample is predicted to have a normal allele with 28-29 CGG repeats and an expanded allele with a full mutation with more than 200 repeats (Adler et al. 2011), which was confirmed by our data. Notably, we also obtained approximately the same number of molecules for the both expanded and normal alleles, despite the nearly 1 kb difference in molecule size. In the expanded allele, we observed a relatively broad size distribution of lengths in individual molecules, ranging from 272 to just over 400 CGG repeats. Instability of the repeats is a hallmark of repeat expansion disorders and this widening of the distribution is a well-documented biological phenomenon (Nolin et al. 1999) that we hope higher resolution sequencing can more precisely evaluate. Genome-wide coverage plots for all Coriell samples are shown in Supplemental Figure 5.

**Figure 5:**
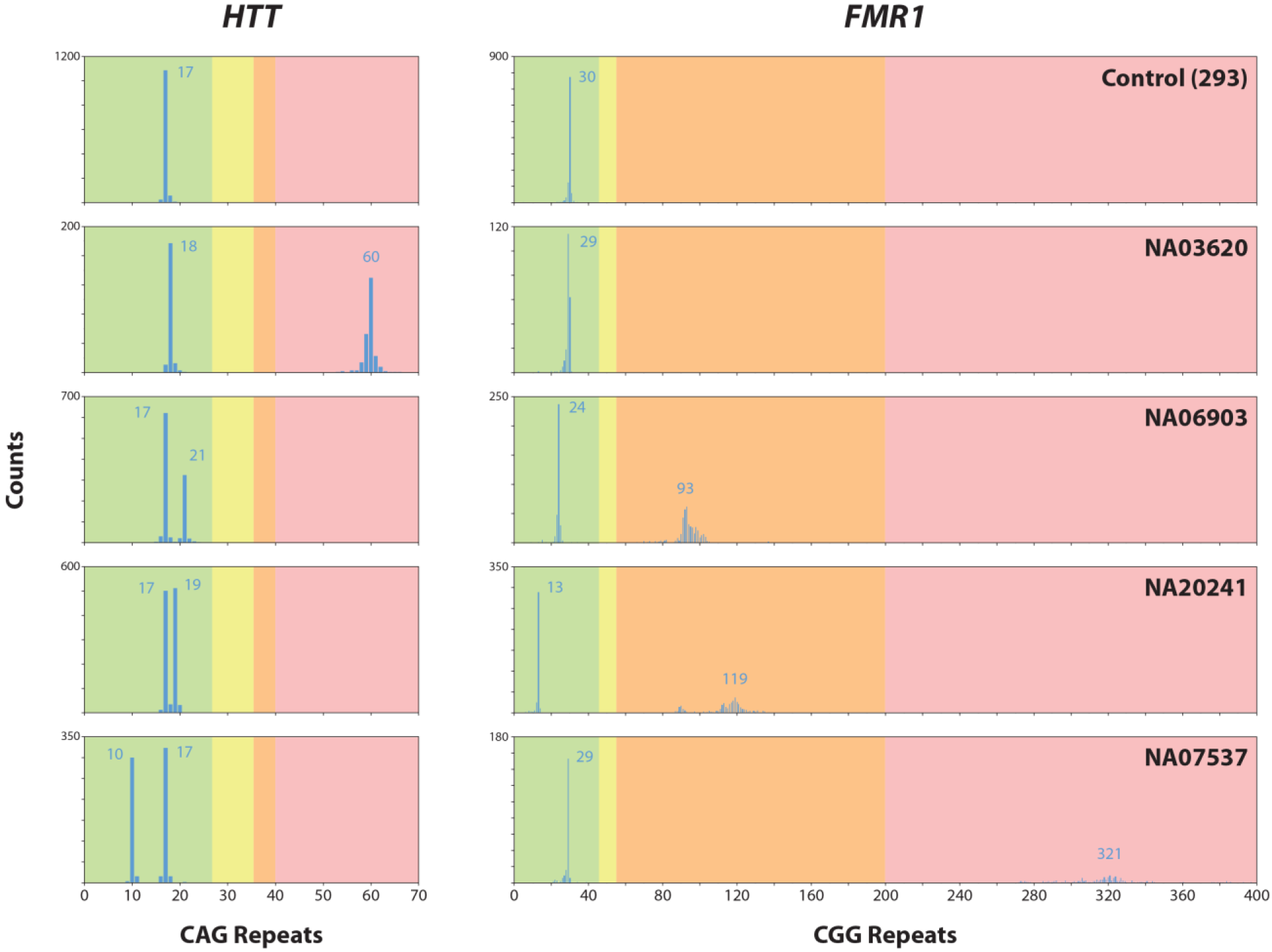
Repeat Count Histograms. Repeat counts are plotted for the *HTT* (left) and *FMR1* (right) loci across all 5 samples with count numbers on the y-axis and CAG (HTT) or CGG (*FMR1*) repeat numbers on the x-axis. Modal values for each allele are labeled. Shaded background in each plot represents risk ranges for developing disease. For *HTT*: green = normal; yellow = high normal (intermediate); orange = incomplete penetrance; red = disease. For *FMR1*: green = normal; yellow = intermediate; orange = pre-mutation; red = full mutation.

## Discussion

The ability to carry out targeted sequencing without the need for amplification unlocks the potential for generating complete sequence information from regions of the genome that cannot be addressed with current NGS approaches. Several of the unique attributes of SMRT sequencing (long reads and unbiased sequencing coverage) make it possible to obtain such sequence information across long repetitive sequences and those containing extreme GC content. The ability to obtain high resolution repeat count information in conjunction with complete and accurate sequence in the same assay is critically important for proper diagnosis of some disorders. The method described here allows for the generation of highly accurate sequencing of hundreds of individual targeted DNA molecules, making it possible to generate distributions of repeat lengths, even for long repeats. Removing the need for PCR amplification that is biased towards shorter alleles is likely to improve the accuracy of measurement of somatic repeat variation present in genomic DNA. Together with the ability to capture both repeat sequence variants (interruptions) as well as extended sequence *in cis* to the repeat, this methodology will facilitate studies aimed at understanding mechanisms of repeat instability

The thousands of reads obtained from a 1 kb genome fragment represents an enrichment factor of approximately 100,000 fold, nearly two orders of magnitude higher than other commonly used enrichment capture methods (Garcia-Garcia et al. 2016). We have also shown that it is possible to target multiple regions of interest in the same reaction. Increasing the multiplex from one to four targets had no adverse impact on the number of molecules captured per locus, suggesting the capacity for targeting an even larger number of regions. Thus, it may be possible to develop large panels of target loci related to a specific research question or field of interest. Furthermore, one could add barcodes to the initial SMRTbell library, thereby enabling multiplexing of patient samples on the same SMRT Cell. This would decrease the input requirements on a per-sample basis, but would result in the division of the total number of reads across the samples. Improvements to the capture method that increase the on-target rate, perhaps by decreasing the random genomic background, will make it possible to multiplex larger numbers of samples in a single SMRT Cell.

In principle, our method could be used for targeted sequencing of nearly any region of the genome or panel of multiple loci. Some regions of the genome may require using different restriction enzymes for the initial preparation of SMRT bell library prior to Cas9 targeting. It is also possible to start by random fragmentation of the genome. For the study of repeat expansions, we chose to employ restriction enzymes to ensure that the captured fragments would contain the entire repeat region. In the process of selection of restriction enzymes and Cas9 guide RNAs, it is important to consider genetic variation. The presence of single nucleotide polymorphisms (SNPs) may disrupt the targeting scheme by introducing or removing a restriction site or decreasing the efficiency of Cas9 digestion. This could be mitigated by using a different combination of restriction enzymes and/or guide RNAs on the same sample in a separate reaction. Separate captures could be combined prior to sequencing.

In this work, we focused on repeat expansions, but the method also has other possible uses. Because we are not employing amplification of any type and sequencing native DNA, it may be possible to exploit another feature of SMRT Sequencing and obtain base modification information across the targeted regions (Flusberg et al. 2010; Clark et al. 2011). DNA methylation has already been implicated in the mechanisms of disease in several repeat expansion disorders (He and Todd 2011). Rather than enriching for loci containing repeat expansions, it should be possible, for example, to target a panel of gene promoters to look at methylation status. The ability to capture large fragments of the genome and generate native sequence from them also permits phasing of disease-associated polymorphisms to normal or expanded alleles. This could have implications for disease prognosis (Becanovic et al. 2015). The method could also serve as a means of testing off-target effects of crRNAs designed for genome engineering in any organism (Hendel et al. 2015).

We have developed a versatile and universal method for target enrichment and sequencing that does not rely on amplification. In conjunction with PacBio SMRT Sequencing, we are able to generate highly accurate consensus sequences for regions of the genome that are difficult to address with other sequencing technologies. In this work, we demonstrate the utility of the method by capturing and sequencing four loci that are associated with repeat expansion disorders. Using patient DNA samples, we sequenced through pathogenic repeat lengths of a CAG repeat from a patient with Huntington’s disease and the CGG repeat from patients with Fragile X/FXTAS. The method also identified allele-specific sequence polymorphisms and repeat interruptions.

## Materials and Methods

### DNA samples used in this study

Human genomic DNA extracted from HEK-293 cells (GenScript) was used for protocol development and protocol optimization experiments. This sample is derived from a female and not known to carry expanded repeats in any of the target loci. Genomic DNA samples carrying previously characterized trinucleotide expansions in the *HTT* locus (NA03620) and *FMR1* locus (NA06903, NA20241, and NA07537) were obtained from the Coriell Institute. Coriell DNA samples were extracted from patient derived lymphoblastoid cell lines. Details are outlined in Table 2.

**Table 2:** DNA Samples Used in This Study.

| ID | Source | Sample Description |
| --- | --- | --- |
| 293 | Genscript | Control DNA extracted from 293 cells (human embryonic kidney)<br>Repeat Status Unknown (Female) |
| NA03620 | Coriell | HTT CAG repeats of 18 and 60 (Female) |
| NA06903 | Coriell | FMR1 CGG repeats of 23 and 95 (Female) |
| NA20241 | Coriell | FMR1 CGG repeats of 29 and 93-110 (Female) |
| NA07537 | Coriell | FMR1 CGG repeats of 28-29 and >200 (Female) |

### Guide RNA design for Cas9 digestion

DNA sequence information from the human genome reference (hg19) was used for designing guide RNAs. Candidate target sequences (20 nt) were selected using the CRISPR RNA configurator available on the Dharmacon website (http://dharmacon.gelifesciences.com/gene-editing/crispr-rna-configurator) with the specificity check option enabled against the human genome. Specific crRNA sequences were manually selected from the candidate list where the target capture region was approximately 1 kb, the 3’ end of the guide RNA was oriented towards the region of interest, and the sequence was determined to have no other perfect matches by BLAST search against the human genome.

#### Guide RNA Sequences

*C9orf72a*: 5’-GCAAUUCCACCAGUCGCUAG-3’
*C9orf72b*: 5’-GCAUGAUCUCCUCGCCGGCA-3’
*FMR1*: 5’-AGAGGCCGAACUGGGAUAAC-3’
*HTT*: 5’-AGCGGGCCCAAACUCACGG U-3’
*ATXN10*: 5’-AUACAAAGGAUCAGAAUCCC-3’

### Input SMRTbell template library preparation

Approximately 5-20 μg of genomic DNA was fragmented with high fidelity restriction enzymes EcoRI-HF and BamHI-HF (New England Biolabs) and SMRTbell template libraries were prepared by ligation of hairpin adapters carrying specific overhang sequences (5’-GATCATCTCTCTCTTTTCCTCCTCCTCCGTTGTTGTTGTTGAGAGAGAT-3’ and 5’-AATTATCTCTCTCTTTTCCTCCTCCTCCGTTGTTGTTGTTGAGAGAGAT-3’) using *E. coli* DNA ligase (New England Biolabs). To improve the observed on-target capture rate, a genome complexity reduction step was added before library preparation by pre-digesting genomic DNA samples with high fidelity restriction enzymes, KpnI-HF, MfeI-HF, SpeI-HF, and EcoRV-HF (New England Biolabs), in the presence of calf intestinal alkaline phosphatase (CIP, New England Biolabs). The four selected nucleases are predicted to have no cut sites within the SMRTbell templates carrying the targets of interest in this study. KpnI-HF and MfeI-HF were used in experiments where only two complexity reduction enzymes were employed.

### Cas9 digestion and asymmetric SMRTbell preparation

The crRNA and tracrRNA were obtained from Integrated DNA Technologies containing an Alt-R modification and annealed at a 1:1 ratio. Up to 1 μg of SMRTbell template library was used in a typical Cas9 nuclease digestion experiment with up to four guide RNAs in the same digestion reaction. Cas9 nuclease from two different vendors (New England Biolabs, Integrated DNA Technologies) were tested and yielded similar results. Following Cas9 digestion, DNA samples were ligated with a polyA hairpin adapter (5’-ATCTCTCTCTTAAAAAAAAAAAAAAAAAAAAAAATTGAGAGAGAT-3’) to obtain asymmetric SMRTbell templates from Cas9-digested target DNA molecules.

### MagBead capture and SMRT sequencing

PacBio MagBeads (part# 100-134-800) were used for enriching the asymmetric SMRTbell templates produced from Cas9 digestion and polyA hairpin ligation. Magbead-DNA binding was carried out in high salt buffer (20 mM TrisHCl pH 7.5, 1 M LiCl, 2 mM EDTA) for 1hr at room temperature and washed once with low salt buffer (10 mM TrisHCl pH 7.5, 0.15 M LiCl, 1 mM EDTA). Bound DNA was then eluted in EB buffer (PacBio Part# 100-159-800) for 10 min at 65 °C.

For SMRT sequencing, a standard PacBio sequencing primer lacking a polyA sequence was annealed to the eluted SMRTbell template (PacBio part# 001-560-849) and purified with 0.6x AMPure beads (PacBio part# 100-265-900) to remove unbound primers. A modified polymerase binding protocol with free hairpin adapters in the binding buffer was used to bind excess DNA polymerase. Sequencing data were collected on a PacBio RSII instrument using the one-cell-per-well MagBead sequencing protocol, P6/C4 sequencing chemistry, and 4-hour collection time.

### Data Processing and Repeat Counting

SMRT sequencing data generated from the asymmetric SMRTbell templates was first subjected to an adapter recall process using a customized PacBio primary analysis software pipeline to identify the polyA hairpin adapter sequence and separate subreads correctly. PacBio SMRTPortal (version 2.3) software was used to generate circular consensus sequencing (CCS) reads and resequencing analysis to provided overall sequencing coverage across the human genome reference (hg19). The length of the repetitive target region was determined by aligning each read against a pool of reference sequences with variable repeat length using blasr (Chaisson and Tesler 2012) and deduced from the reference sequence which provided the best overall alignment score.

## Supplemental Material

Supplemental Figure 1: Cas9 Target Loci Schematics

Supplemental Figure 2: Sequencing Statistics

Supplemental Figure 3: Cas9 Off-Target Sites

Supplemental Figure 4: Per Molecule Consensus Accuracy

Supplemental Figure 5: Genome-wide Coverage Plots

Supplemental Table 1: Complexity Reduction Library and Sequencing Yield

## Acknowledgements

The authors would like to thank the Jennifer Doudna lab for initial advice on Cas9 and Ting Hon, Matthew Boitano, Paul Kotturi, Jenny Ekholm, Michael Weiand, and Nicola Cahill for input on the method.

